# Adaptive Evolution for Increased Biomass and Magnetic Nanoparticle Productivity of Magnetotactic Bacteria

**DOI:** 10.1101/022590

**Authors:** Karthik Sekar, Javin P. Oza

## Abstract

Large magnetic nanoparticles (over 25 nm diameter) are a valuable commodity but remain difficult to synthesize using traditional chemical synthesis methods. Magnetotactic bacteria (MTBs) have evolved mechanisms to produce monodisperse, protected magnetic nanostructures within organelles (magnetosomes). Genomic diversity of MTB species result in unique particle properties that vary in shape and size ranging from 30 nm to 150 nm based on the genetic background. Culturing and engineering MTBs for the production of magnetic nanoparticles carries tremendous potential but is underdeveloped. This primarily because MTBs are difficult to culture and genetically manipulate, limitations that could be alleviated with adaptive evolution. We propose the magnetotrophic reactor, a novel bioreactor system for adaptively evolving MTBs for better growth and magnetosome production. This platform is projected to be superior to the traditional evolution methods since robust growth phenotypes can be selected for while maintaining selective pressure for magnetotaxis. We provide, herein, a quantitative basis for our platform including considerations of continuous evolution, sizing, and magnetic field pulsing. Our proposed fermentation process anticipates scalable production of 1 g/L per day of monodisperse magnetic nanoparticles to enable industrial applications.

## 1. Introduction

Magnetic nanoparticles have a variety of uses including catalysis, magnetic fluids, magnetic resonance imaging, data storage, and remediation [1]. Beneath a threshold size, each magnetic nanoparticle is its own magnetic domain; thus, each particle can have a magnetic moment and act as a paramagnetic atom. Synthesis of magnetic nanoparticles, however, is complicated by their reactivity and tendency to aggregate. Under ambient conditions, particles are oxidized and lose their magnetism. Expensive surfactants and reducing agents are required for aqueous-based synthesis methods. Furthermore, traditional methods, such as co-precipitation, tend to result in polydispersed particles complicating downstream application where monodispersity is required[1]. Synthesis methods using organic solvents such as thermal decomposition require organometallic precursors under high temperature conditions to achieve monodispersity and gram per batch quantities[2]. Traditional synthesis methods, however, tend to achieve a maximum particle size of 25 nm, which is too small for applications such as magneto-hypertheramic treatment of tumors [3].

Magnetotactic bacteria (MTBs), native to stratified layers at the bottom of the ocean, naturally produce monodisperse magnetic nanoparticles such as magnetite (Fe_3_O_4_) at size ranges typically inaccessible from traditional synthesis methods (30 to 150 nm diameter). They use the magnetic nanoparticles to passively align to the Earth’s magnetic field so that their search is one-dimensional at a given oxygen concentration (which varies with depth in aquatic environments). Particle size are monodisperse within a given organism and vary in size and shape between species [4]. In order to make such nanoparticles, MTBs synthesize the magnetic crystal within an organelle, the magnetosome, where a phospholipid layer covers the crystal. Individual magnetosomes are secured to a cytoskeletal filament that runs the length of the organism, thus allowing the organism to align to Earth’s magnetic field. The magnetic nanoparticles are biosynthesized within the periplasm. Proteins directly involved have been characterized for their role in the process. The primary steps in the biosynthesis start with the invagination of the inner membrane. The cavity formed will be the nest for the growing nanoparticle. Membrane associated magnetosome proteins then get sorted and positioned through the inner membrane. This process happens in parallel such that several invagination events occur side-by-side in a chain-like orientation. Iron is then delivered into the cavity and crystal formation begins. Size and shape of the crystal are tightly controlled as the crystal matures[5].

Making magnetite via the production of magnetosomes is an attractive avenue. However, MTBs are notoriously difficult to culture, and much research has been dedicated to heterologously expressing magnetosome-forming genes in other organisms. The Schuler group has successfully imported machinery from the MTB, *M. gryphiswaldense* to the phylogenetically similar *R. rubrum* but with smaller sizes and inconsistent shapes [6]. The Lee group has engineered *E. coli* to make metal nanoparticles but with much smaller size ranges compared to that of MTBs [7]. The Silver group has engineered yeast to show a magnetic attraction phenotype; however, their yeast did not synthesize particles [8]. Personal discussions with Schuler suggest that importing magnetosome machinery in to production hosts such as *E. coli* and *S. cerevisae* is ongoing but difficult. We propose, instead, to directly engineer and evolve magnetotactic bacteria toward the easier culture conditions and increased biomass and magnetite production.

MTBs cultures have poor biomass yields; generally the total biomass in the state of art fermentation is an optical density (OD) of 12 (or 3 g biomass/L) after 40 hours [9]. In comparison, an *E. coli* culture can reach an OD of 100 (or 40 g biomass/L) in half the time [10]. There have only been a few studies designing optimal culture conditions for MTBs. A continuous fermentation was able to produce 55 mg of ferrites over one day per liter [9], which is up from 17 mg/L/day of ferrite six years prior [11]. We see MTBs themselves as a potentially productive host given the proper selection and genetic engineering. In particular, our adaptive evolution techniques, unique to MTBs, select for strains that adapt to cheaper media, dissolved oxygen, and ones that grow faster with increased yield all while maintaining pressure to continue producing magnetic nanoparticles. We expect to approach 1 g/L/day of crystal with our strategy based on metrics from previous adaptive evolution efforts.

## 2. Reactor Basics and Assumptions

Traditional methods for selection of high biomass producing strains such as serial shake flask dilution cannot be used because variants may produce magnetite crystals less productively [12]. The proposed reactor, the magnetotrophic bioreactor (Figure 1), is a long, cylindrical reactor where a MTB culture is seeded on one end and a magnetic field runs across leading bacteria to the other end with more nutrients. The magnetotrophic reactor is designed to provide selection toward both growth and magnetosome productivity.

**Figure 1.**
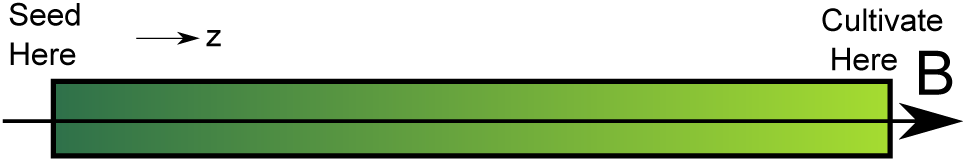
The magnetotrophic reactor selects for both growth and magnetite productivity. The basic design for the reactor is analogous to a plug flow system (but with no bulk fluid flow) with an applied magnetic field *B* running axially. Microbes are seeded at the terminal seed end (left end) and cultivated from the right end. The *z* direction is axial to the reactor.

We make a number of key assumptions including as follows:

- Minimal mixing.
- Motility of MTBs >> Diffusion of MTBs
- Minimal biofilm formation for continuous conditions
- Variance only along the z axis. No variation radially.
- There exists a distribution of speed and magnetosome productivity among a population of MTBs.

### 2.1 Definitions

*X* - cell concentration (cells/vol)

*μ* - the growth rate (1/hr)

*V*_*total*_ - reactor volume (vol)

*Q* - bulk movement rate (vol/hr)

*v* - the average velocity of each cell (50 *μ*m/s)

*S* - substrate concentration (mol/vol)

*D* - dilution rate (1/hr)

*Y*_*X*/*S*_ - biomass yield on substrate (cells/mol)

*m* - magnetic moment of each MTB(5 × 10^−17^*Am*^2^) [13]

*B* - magnetic field

## 3. Exponential Configuration

We first start with an exponential configuration that is analogous to serial dilution of culture in shake flasks (Figure 2). To understand this setup, we derive the characteristic design equations. Assume that the reactor is fully perfused with nutrients and media.

**Figure 2.**
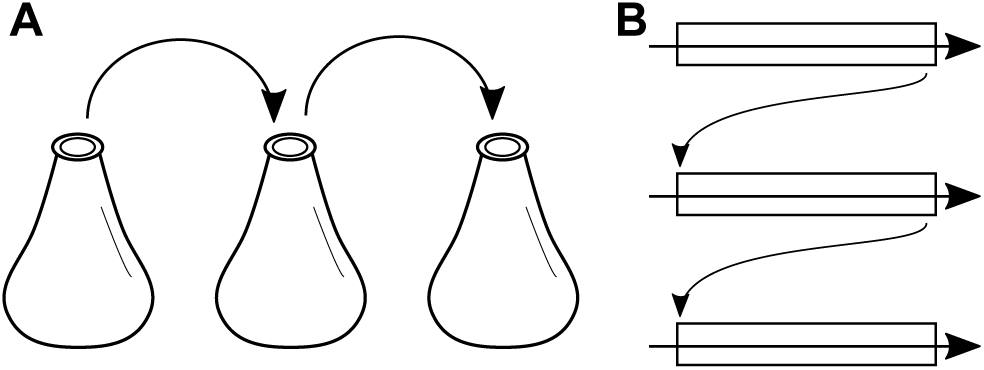
Exponential growth magnetotrophic reactor configuration. Typical growth-based selection may occur with serial shake flask dilution (**A**). Flasks are seeded, grown, and then diluted into fresh shake flasks to start a consequent cultivation. This exponential configuration, where nutrients are not limiting, can be achieved with the magnetotrophic reactor system by seeding with culture from the terminal (right) end of a previous round (**B**).

Applying a chemical balance to a differential volume within the reactor:

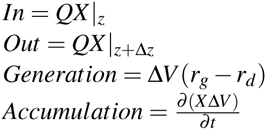

Assume that the death rate (*r*_*d*_) is negligible and there is no nutrient limitation; therefore, *r*_*g*_ = *μX*. This yields

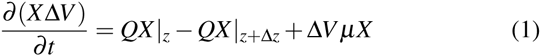

Rearrange and simplify to

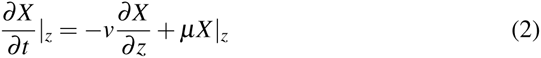

The resulting equation describes the how the biomass changes over time in response to two mechanisms: motility and growth. In contrast, a shake flask would simply have the growth term, *μX*.

## 4. Continuous Growth Configuration

For the continuous system, our process is analogous to a chemostat as shown in Figure 3. In this configuration, three flow streams are introduced as a function of an assigned flow rate *F*. An in stream on the right end of the reactor delivers fresh, sterile media. An outlet, waste stream on the left removes culture. A recycle stream (< *F*) continuously reseeds the reactor. The specific flow rate for the recycle stream may need to be optimized but should be less than the inlet and outlet streams. Note that despite the directionality of the flow, there is not supposed to be bulk flow in the −*z* direction. The flow is simply for media replenishment and culture outflow. As with a chemostat, the design equation of relating the dilution rate *D* to the flow rate *F* and reactor volume *V*_*total*_ still applies where 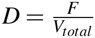.

**Figure 3.**
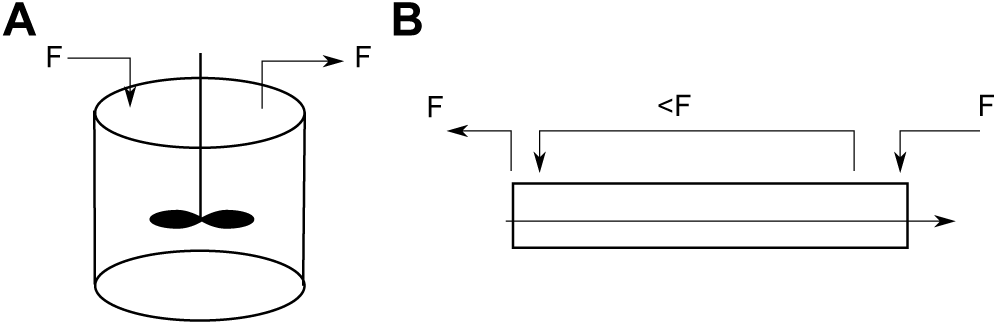
Continuous growth magnetotrophic reactor configuration. A chemostat is a steady-state bioreactor where an inlet stream delivers fresh, sterile media and an outlet stream removes growing culture (**A**). Chemostats provide continuous evolution and select for strains with the best biomass yield. An analogous set up for magnetotrophic reactor is achievable (**B**). A recycle stream is added to reseed best performing MTBs from the cultivation end back to the seed end. The recycle flow rate must be less than the overall flow rate for the system.

The reactor is substrate limited along *z* axis, *S*(*z*). Equations remain the same as the exponential configuration except the rate of generation has changed to Δ*Vr_g_* = *D*Δ*VY*_*X*/*S*_*S*(*z*). With the replaced term:

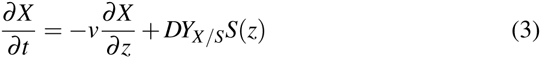

In steady state configuration, by imposing chemostat configuration over entire reactor (the flow rate *F*):

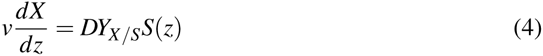

The resulting equations suggests that the dilution rate *D* must be tightly controlled otherwise there will be poor selection (too low of a *D*) or wash out (too high of *D*). This system can be combined with an online turbidity measurement to adjust *D* automatically relative to the aggregate growth rate *μ* of all the cells collectively (Figure 4). In such a configuration, the system would toggle between two states. In the *D* < *μ* state, this system would experience exponential growth because cells are growing faster than being removed. In the opposite state, *D* > *μ*, the system would experience washout particularly of the strains that grow and/or move slowly. This automated system may provide a less laborious means to adaptively evolve strains.

**Figure 4.**
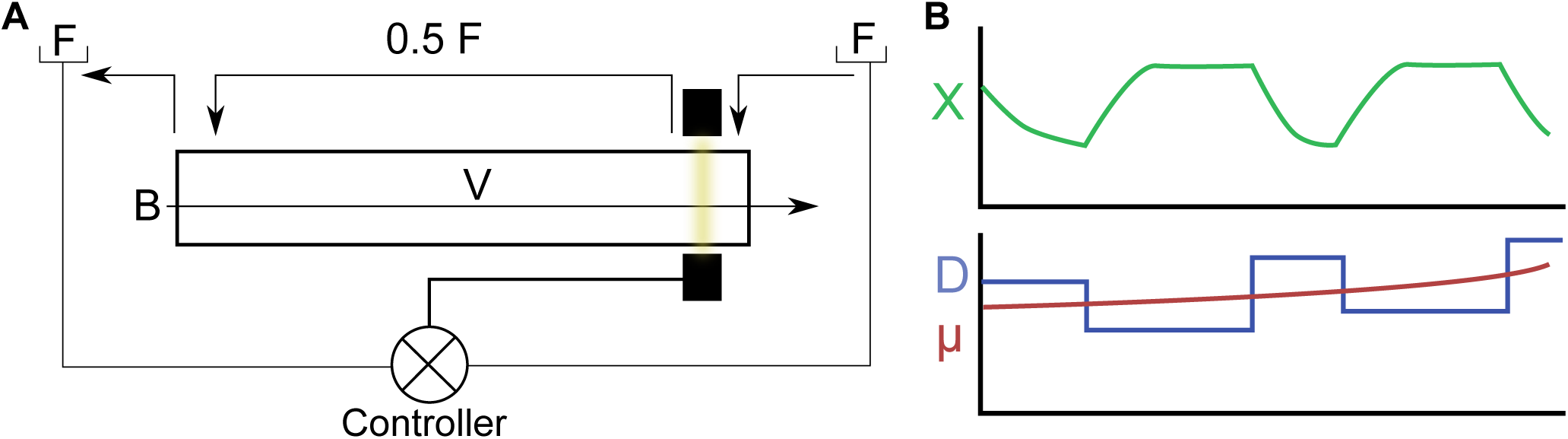
Online turbidity measurement can be used to provide continuous evolution (**A**). The turbidity measurement in conjunction with a controller can be used to modulate the dilution rate. The dilution rate can be pulsed higher and lower than the aggregate growth rate for the system (**B**). When *D > μ* then the slow growers and movers are washed out of the system, when *D* < *μ*, the system allows the culture to reach steady state again. The turbidity measure can be used to switch the dilution rate accordingly.

## 5. Relative Selection Pressures

We can roughly calculate the relative selection pressure between growth and motility by comparing the orders of magnitude of our two terms. To simplify calculations, normalized cellular concentrations are used instead, 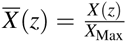. To have comparable selection from each, for example, we desire the magnitudes to be roughly equal:

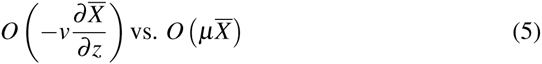

If we use approximate numbers and make some assumptions (reactor length is 10 cm), we find:

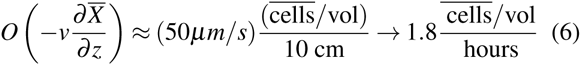

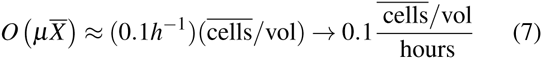

In this case, the selection pressure of motility is approximately 18 times greater than growth. Note that simply by lengthening the reactor, selection can be tilted toward growth. For example, using a reactor with length of approximately 180 cm would equalize the selection pressures.

## 6. Magnetic Field Strength and Pulsing

The magnetic field of the Earth is very weak (around 0.5 gauss). This would suggest that any applied magnetic field would immediately align the MTBs and not necessarily incentivize them to make magnetosomes productively. This problem can be overcome by pulsing the magnetic field. By pulsing, only the MTBs with higher magnetic moments will align quickly (Figure 5). Higher magnetic moments by individual MTBs can be achieved by producing more magnetosomes. Optimal pulsing would need to be determined through experimental measurements and possibly modeling techniques (e.g. agents-based).

**Figure 5.**
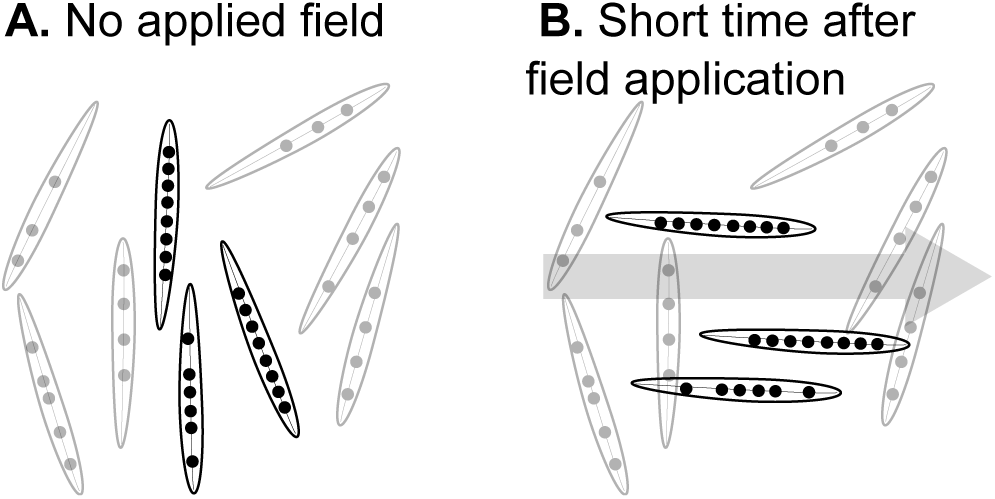
Pulsing the magnetic field can be used to select for the strains that produce magnetosomes more productively. In the absence of no magnetic field, MTBs are aligned to the Earth’s magnetic field or randomly ordered (**A**). When the magnetic field *B* is pulsed, the strains with higher magnetic moment (more magnetosome chains) will align faster to the field (**B**). Pulsing features can be integrated into the magnetotrophic reactor system.

Considering Earth’s magnetic field (*B*_*earth*_ = 0.5 gauss) and the magnetic moment of each MTB (*m* = 5 × 10^−17^*Am*^2^), we can calculate the maximum needed torque, *T*_*max*_ = *B*_*earth*_*m* = 2.5 × 10^−21^*Nm*, for the alignment of MTBs. This torque is low; therefore, when pursuing the pulsing configuration, the maximum quantity of magnetic field and frequency need to be carefully modulated and can be constrained by the following equation:

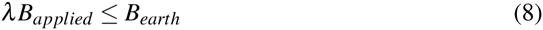

Where *λ* is the ratio of magnetic field time ON versus time ON + time OFF, and *B*_*applied*_ is the strength of the applied magnetic field. Pulsing would also likely change the aggregate speed, *v*, of the MTBs through the reactor. The new speed *v* would be between *λv_max_* and *v*_*max*_. This would additionally allow for smaller reactor sizes.

## 7. Outlook and Discussion

To evaluate the potential metrics of evolved strains, we consider previous adaptive evolution efforts. To our knowledge, no adaptive evolution cultures have been performed with MTBs. In comparison, there have been adaptive evolution studies with *E. coli* and *S. cerevisae*. Yeast (e.g. *S. cerevisae*) have more comparable doubling times to MTBs (around 2 hours for yeast versus 5 to 10 for MTBs). One recent study was able to increase yeast growth rate almost two fold in high temperature conditions and the biomass yield 20% [14]. Similar projections for MTBs would suggest fermentation times decreasing from 40 hours (7 doublings) to 20 hours with higher cell density. However, since MTBs are typically from nutrient poor conditions, the most representative case may be an adaptive evolution of *Geobacter sulfurreducens* where the iron uptake was increased 1000% [15].

Desensitizing MTBs to oxygen may help downstream bioreactor scale up. Most species of MTBs grow under suboxic conditions and tend not to make magnetite or grow well under oxygenated or completely anaerobic conditions [16]. Decoupling this phenomenon may increase biomass yields and growth rates because fermentation oxygen concentration may not be able to be controlled so precisely. Note that oxygen has a higher redox potential than iron. Oxygen may be competing with ferrite for electrons and the amount oxygen may need to be controlled in the reactor in a manner that adaptive evolution may not be able to solve.

To save on media costs and to develop strains to grow in defined, minimal media conditions, we propose to select for strains to tolerate and thrive on the cost-saving minimal media. Highest biomass yield on media was achieved using an undefined, rich media [11]. This selection can be performed by slowly changing the composition of the magnetotrophic reactor over time from the rich to progressively more minimal media. Particular media components to replace include yeast extract with defined amino acids. Additionally, the media can contain lactic acid, a fermentation product of MTBs that also inhibits their growth[9]. Strains that grow better on lactic acid would be more amenable to downstream fermentation applications.

Lastly, this system complements with mutagenesis and genetic engineerings strategies designed to create a large number of candidate strains. A population with more variable magnetosome productivity and growth rates can be fed into the magnetotrophic reactor to enrich the pool for either the exponential or continuous configuration. This may reduce the number of generations needed to achieve a strain with the desired features.

## 8. Conclusions

We have proposed an adaptive evolution scheme for MTBs to select for magnetosome productivity and growth. Our platform is strain agnostic and low risk technically. We have derived the design equations associated with the magnetotrophic reactor that help with sizing and pulsing the magnetic field. We show that this device can easily be developed physically for example using PVC piping with a live wire wrapped around to form a solenoid.

## Acknowledgments

We acknowledge Stephanie R. Jones (University of California at Berkeley), Keith E.J. Tyo (Northwestern University), and Wesley Burghardt (Northwestern University) for helpful discussions and feedback. We also thank Rajesh Naik (Air Force Research Laboratory) and Andrew Herr (Helicase and Mind Plus Matter) for choosing a proposal based on this work for the ‘Synthetic Biology for Materials’ InnoCentive challenge.

